# Phase of Firing Coding of Learning Variables across Prefrontal Cortex, Anterior Cingulate Cortex and Striatum during Feature Learning

**DOI:** 10.1101/2019.12.12.874859

**Authors:** Benjamin Voloh, Mariann Oemisch, Thilo Womelsdorf

## Abstract

The prefrontal cortex and striatum form a recurrent network whose spiking activity encodes multiple types of learning-relevant information. This spike-encoded information is evident in average firing rates, but finer temporal coding might allow multiplexing and enhanced readout across the connected the network. We tested this hypothesis in the fronto-striatal network of nonhuman primates during reversal learning of feature values. We found that neurons encoding current choice outcomes, outcome prediction errors, and outcome history in their firing rates also carried significant information in their phase-of-firing at a 10-25 Hz beta frequency at which they synchronized across lateral prefrontal cortex, anterior cingulate cortex and striatum. The phase-of-firing code exceeded information that could be obtained from firing rates alone, was strong for inter-areal connections, and multiplexed information at three different phases of the beta cycle that were offset from the preferred spiking phase of neurons. Taken together, these findings document the multiplexing of three different types of information in the phase-of-firing at an interareally shared beta oscillation frequency during goal-directed behavior.

**Highlights:**

- Lateral prefrontal cortex, anterior cingulate cortex and striatum show phase-of-firing encoding for outcome, outcome history and reward prediction errors.
- Neurons with phase-of-firing code synchronize long-range at 10-25 Hz.
- Spike phases encoding reward prediction errors deviate from preferred synchronization phases.
- Anterior cingulate cortex neurons show strongest long-range effects.

## Introduction

The lateral prefrontal cortex (LPFC) and anterior cingulate cortex (ACC) are key brain regions for adjusting to changing environmental task demands (Murray and Rudebeck, 2018; White et al., 2019). Both regions project to partly overlapping regions in the anterior striatum (STR), which feeds back projections and thereby closes recurrent fronto-striatal loops (Haber and Knutson, 2010). Neurons in this recurrent network encode multiple learning variables during goal-directed behaviors including the value of currently received outcomes, a memory of recently experienced outcomes, and a reward prediction error that indicates how unexpected currently received outcomes were given prior experiences (Hikosaka et al., 2019; Oemisch et al., 2019).

The multiplexing of outcomes, outcome history, and outcome unexpectedness (prediction errors) within the same neuronal population is likely realized by firing rate modulations in fronto-striatal brain areas (Fusi et al., 2016), but how this firing is temporally organized within the larger network is unresolved (Akam and Kullmann, 2014; Kumar et al., 2010; Panzeri et al., 2015). A large body of evidence has shown that ACC and LPFC synchronize their local activities at a characteristic beta oscillation frequency (Babapoor-Farrokhran et al., 2017; Nácher et al., 2019; Voloh and Womelsdorf, 2017a; Womelsdorf et al., 2014a, 2014b), and that both areas engage in transient beta rhythmic oscillatory activity with the striatum during complex goal directed tasks (Antzoulatos and Miller, 2014; Feingold et al., 2015; Howe et al., 2011; Leventhal et al., 2012). However, whether this beta oscillatory activity is informative for learning and behavioral adjustment has remained unresolved (Amemori et al., 2018; Spitzer and Haegens, 2017; Womelsdorf and Everling, 2015). Prior studies have documented that beta activity emerges specifically during the processing of outcomes following correct trials during habit learning (Howe et al., 2011), and that following error trials overall beta activity is larger when the committed error is smaller (Tan et al., 2014a). However, these studies did not quantify whether neuronal spiking activity synchronizing to these beta oscillations contains learning relevant outcome information.

We therefore aimed to clarify how outcome related beta rhythmic activity relates to the behavioral learning of reward rules in ACC, LPFC, and STR. First, we quantify firing rate information about current outcomes, about prediction errors of these outcomes and the history of recent reward. These variables might be conveyed independently of network level beta oscillatory activity. However, theoretical studies suggest that neuronal coding utilizing temporal organization can be efficient, high in capacity, and robust to noise (Akam and Kullmann, 2014; Kayser et al., 2009; Kopell et al., 2011; Kumar et al., 2010). Additionally, coding of information in the temporal activity pattern has been linked to mechanisms of efficient communication among neuronal groups, suggesting that coherently synchronized groups can exchange information by phase aligning their disinhibited activity periods (Fries, 2015; Hahn et al., 2014, 2019; Luczak et al., 2015; Palmigiano et al., 2017; Singer and Gray, 1995; Womelsdorf and Fries, 2007).

To this end, we recorded from LPFC, ACC, and STR while macaque monkeys engaged in trial-and-error reversal learning of feature reward rules. We found that during outcome processing, each area contained segregated ensembles of neurons encoding the current *Outcome*, the *Prediction Error* of those outcomes, and the recent *Outcome History* in their overall firing rates. A large proportion of rate coding neurons synchronized long-range to remote areas of the fronto-striatal network at a shared 10-25 Hz beta frequency range. We found that for those neurons that phase synchronized long-range, the three learning variables were encoded more precisely for spike output elicited at narrow oscillation phases in the beta band. This phase-of-firing gain of encoding exceeded the firing rate code and occurred at phases that were partly away from the neurons preferred spike phase. These findings document a multiplexing of learning variables through the phase of firing across the ACC, LPFC, and STR of nonhuman primates.

## Results

Animals performed a feature-based reversal learning task (Oemisch et al., 2019). Subjects were shown two stimuli with opposite colors and had to learn which of them led to reward (**Figure 1A**). The same color remained associated with reward for at least 30 trials before an uncued reversal switched the color-reward association (**Figure 1B**). During each trial, the subjects monitored the stimuli for transient dimming events in the colored stimuli. They received a fluid reward when making a saccade in response to the dimming of the stimulus with the reward associated color, while the dimming of the non-reward associated color had to be ignored. A correct, rewarded saccade to the dimming of the rewarded stimulus had to be made in the up- or downward direction of motion of that stimulus. This task required covert selective attention to one of two peripheral stimuli based on color, while the overt choice was based on the motion direction of the covertly attended stimulus. In 110 and 51 sessions from monkey’s HA and KE, respectively, we found that subjects attained plateau performance of on average 80.2% (HA: 78.8%, KE: 83.6%) within 10 trials after color reward reversal (**Figure 1C**). Using a binomial General Linear Model (GLM) to predict current choice outcomes (correct or error outcomes, excluding fixation breaks), we found that for both subjects, outcomes from up to three trials into the past significantly predicted the current choice’s outcome (**Figure 1D**; Wilcoxon signrank test, p<0.05, multiple comparison corrected), closely matching previous findings (Walton et al., 2010).

**Figure 1.**
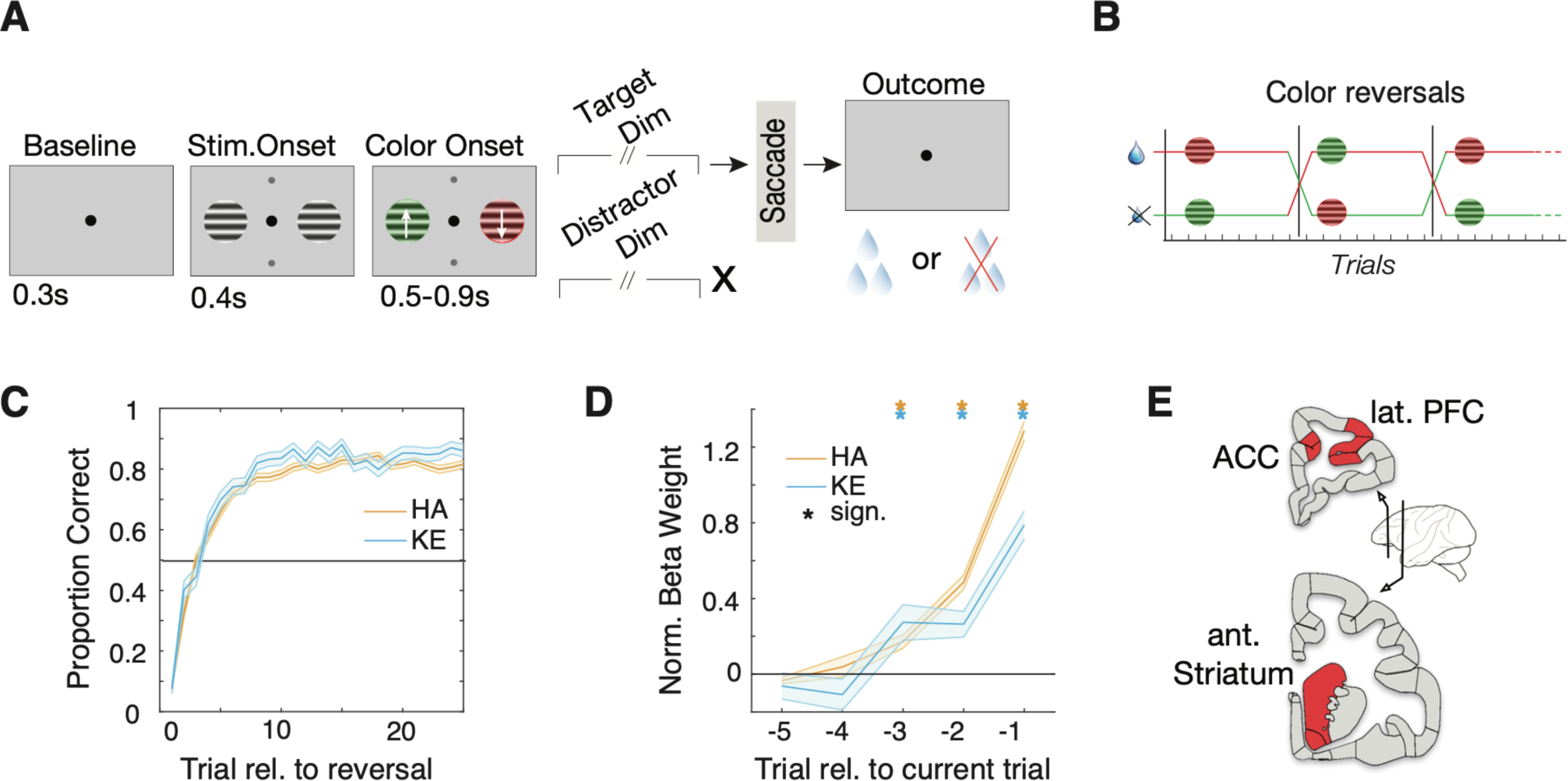
Subjects learned reversals. **(A)** Feature-based reversal learning task. Animals are presented with two black/white stimulus gratings to the left and right of a central fixation point. The stimulus gratings then become colored and start moving in opposite directions. Dimming of the stimuli served as a Go signal. At the time of the dimming of the target stimulus the animals had to indicate the motion direction of the target stimulus by making a corresponding up or downward saccade in order to receive a liquid reward. Dimming of the target stimulus occurred either before, after or at the same time as the dimming of the distractor stimulus. **(B)** The task is a deterministic reversal learning task, whereby only one color is rewarded in a block. This reward contingency switches repeatedly and unannounced in a block-design fashion **(C)** Accuracy relative to block start for monkey HA (orange) and KE (blue). Shaded region represents the standard error. Subjects achieved plateau performance within 5-10 trials **(D)** Median beta coefficients from a binomial regression of current outcome as predicted by past outcomes. Shaded region represents the standard error. Outcomes up to three trials into the past predicted current outcome (Wilcoxon signrank. p<0.05, multiple comparison corrected). **(E)** Schematic depicting recorded brain areas.

### ACC, LPFC and STR Neurons Encode Outcomes, their History and their Prediction Error

To test how previous and current outcomes are encoded at the single neuron level, we analyzed a total of 1460 neurons, with 332/227 (monkey HA / KE) neurons in lateral prefrontal cortex (LPFC), 268/182 neurons in anterior cingulate cortex (ACC), and 221/230 neurons in anterior striatum (STR) (**Figure 1E, Figure S1A**). These regions have previously been shown to encode outcome, outcome history, and prediction error information (Bernacchia et al., 2011; Hikosaka et al., 2018; Ito, 2003; Lee et al., 2012; Oemisch et al., 2019; Shen et al., 2014; Yamada et al., 2011). We found multiple example neurons encoding different types of outcome variables. Some cells responded differently to correct versus erroneous trial outcomes irrespective of previous outcomes (**Figure 2A**), while others responded strongest when the current outcome deviated from the previous trials’ outcome (signalling reward prediction error) (**Figure 2B**), or when the current outcome was similar to the previous trials’ outcome, i.e. following a sequence of correct trials or a sequence of error trials (**Figure 2C**).

**Figure 2.**
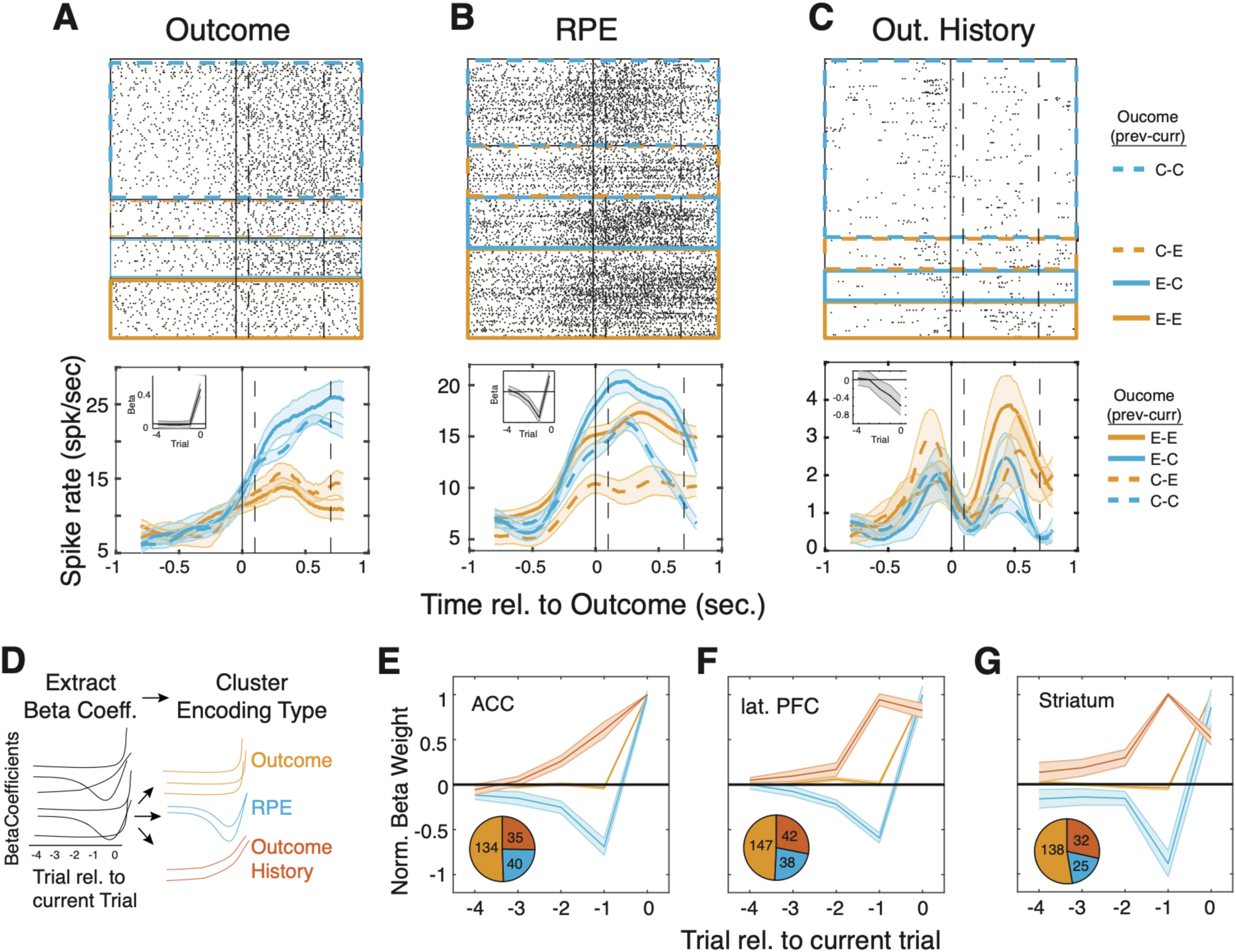
Outcomes, prediction errors, and outcome history is encoded across the fronto-striatal axis. **(A-C)** Examples of *Outcome (A), RPE (B)*, and *Outcome History (C)* cells. (*top)* Raster plot for four separate trial conditions. Color denotes outcome on the current trial (blue=correct, yellow=incorrect), and line style to the previous trial (solid=error, dashed=correct). The dotted vertical lines represent the time period used for the GLM analysis. (*bottom*) Time-resolved firing rate for the four different trial types. Insets in the top left depict the average beta coefficients. **(D)** Sketch of general approach. We regressed spike counts *C* in the [0.1 0.7] outcome period against outcomes up to 5 trials into the past using a penalized LASSO GLM. Beta coefficients were then clustered to group cells according to their most parsimonious functional designation. **(E-G)** Median normalized beta coefficients for three functional clusters in ACC (E), LPFC (F), and STR (G). Shaded regions represent the standard error. On the basis of the pattern of beta weights (see text), each area exhibits “*Outcome*” cells (yellow), “*Outcome History*” cells (red), and “*RPE*” cells (blue). *(inset)* Relative frequency of each functional cluster. Outcome cells were the most populous in all three regions (Chi-squared test, p<<0.05).

We quantified these types of outcome encoding using a LASSO Poisson GLM that predicted the spike counts during the outcome period (0.1-0.7 sec after reward onset) and extracted the characteristic patterns of beta weights across the past and current trial outcomes that distinguished different types of outcome encoding (**Figure 2D**). Neurons that encoded only the current trials’ outcome showed large weights only for the current trial (*Outcome* encoding type). Neurons encoding a prediction error showed beta weights for previous trials that were opposite in sign to the current trial’s outcome (*Prediction Error* encoding type). In neurons encoding the history of recent rewards, beta weights ramped up over recent trials toward the current trial outcome (*Outcome History* encoding type) (*see* also insets in **Figure 2A-C**). We found in a clustering analysis that these three types of outcome encoding were prevalent in each of the recorded brain areas (**Figure 2E-G, Figure S1B**). Neurons encoding the current outcome were the most populous (∼59%, 234/384 in monkey HA and 185/329 in monkey KE), followed by ∼26% of neurons encoding *Prediction Errors* (64/231 in monkey HA and 39/170 in monkey KE) and ∼32% of neurons encoding *Outcome History* (76/206 in monkey HA and 33/139 in monkey KE) (**Figure 2E-G**; χ^2^ test, p<0.05). The relative frequency of these encoding types did not differ between areas (χ^2^ test, p=0.46). Across the population of neurons in ACC, LPFC and STR we found that *Outcome, Prediction Error* and *Outcome History* encoding emerged shortly after outcomes were received (see **Methods, Figure S1C,D**, Wilcoxon signrank test, p<<0.05). Neurons encoding *Outcome, Prediction Error*, or *Outcome History* showed similar overall firing rates (**Figure S1E;** Kruskal Wallis test, p_LPFC_=0.27, p_ACC_=0.58, p_STR_=0.35).

### Neuronal Synchronization at 10-25 Hz Beta Band across ACC, LPFC and STR

We found similar proportions and activation time courses of encoding neurons in ACC, LPFC and STR (**Figure S1C,D**), which raised the question how these neuronal populations are connected. One possibility is that neuronal firing patterns are organized temporally, such that spikes in one area phase synchronize to neuronal population activity in remote areas. We assessed the degree of phase consistency of neuronal spikes with local field potential (LFP) fluctuations in distally recorded areas using the Pairwise-Phase Consistency metric (PPC) (**Figure 3A**; (Vinck et al., 2011; Womelsdorf et al., 2010); *see* **Methods**). Across all (n=7938) spike-LFP pairs, we found a pronounced peak of phase synchronization in the beta band (10-25 Hz) with neurons firing on average ∼1.15 times more spikes on their preferred (average) phase than at the opposite phase (**Figure 3B**). Neurons that encoded outcome variables in their firing rates showed similar phase synchrony as neurons not encoding reward outcome information (**Figure 3B**). Across all pairs, 55% (4320/7938) showed significant phase synchronization within the 10-25 Hz range (**Figure 3C;** Rayleigh test, p<0.05, *see* Methods for prominence criteria). Among those neurons that showed significant firing rate encoding of *Outcome, Prediction Error* or *Outcome History*, the most prevalent within-area beta synchrony was evident within the ACC, compared to LPFC (One-way ANOVA, p=0.014) or Striatum (p∼0) (**Figure 3D,E**). Neurons in ACC were also more likely than expected by chance to show between-area spike-LFP synchrony, as compared to LPFC (p∼0) and STR (p=0.042). There was also a trend for stronger between-area synchrony with spikes originating in STR, as compared to LPFC (p=0.058) (**Figure 3D,F**). Testing for the reciprocity of beta band phase synchrony showed that ACC spikes phase synchronized more strongly to beta in the LPFC than vice versa (p=0.047) (**Figure 3G**). Intra-areal LPFC and STR pairs showed statistically indistinguishable spike-phase synchrony strength (p=0.92), as did ACC and STR pairs (p=0.26). Similar findings were evident for the strength of inter-areal spike-LFP phase synchrony (**Figure S2**). Both monkeys showed stronger synchronization for spikes originating in ACC compared to LPFC and STR spike output (**Figure S3B**). These results show that neurons with firing rate information about *Outcome, Prediction Error*, or *Outcome History* synchronized their firing within and between areas at a characteristic 10-25 Hz frequency range.

**Figure 3.**
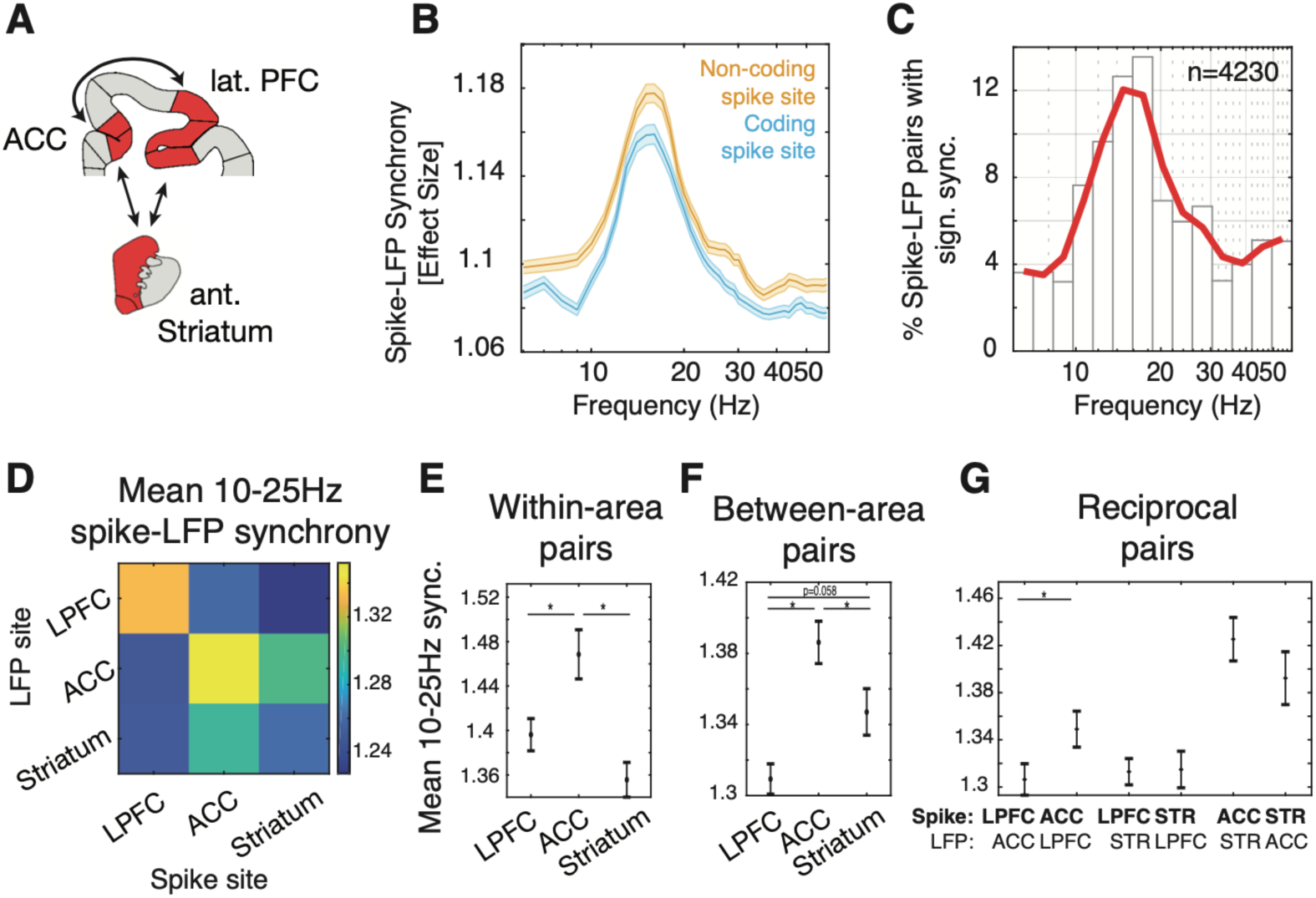
Spike-phase synchronization in beta band tends to occur in ACC. **(A)** We tested for functional connectivity between the ACC, LPFC, and STR. **(B)** Average PPC effect size between distal LFP and spikes at encoding (blue) and non-encoding (orange) sites. Shaded area represents the standard error. There is a prominent beta peak for both. X-axis is depicted on a log-scale for clarity. **(C**) Proportion of spike-LFP pairs that exhibited significant, prominent locking (*see* Methods). **(D)** Average inter-areal synchrony between all pairs of areas for signification, prominent beta in the [10-25] Hz range. Color denotes the average. **(E)** Mean and standard error contrasting spike-phase locking within different areas (diagonal in Fig 2D). ACC synchronizes more strongly that either LPFC or STR. **(F)** Same as (E) but for LFPs originating in other areas (i.e. summing across columns, less the diagonals, in Fig 2D). ACC synchronizes more strongly to distal beta, compared to LPFC or STR. **(G)** All pairwise comparison between regions. ACC spikes lock more strongly to LPFC beta than the inverse.

### Phase-of-Firing at 10-25 Hz Encodes Outcome, Prediction Error and Outcome History

Neurons that synchronized to the LFP elicit more spikes at their mean spike-LFP phase, which we denote as the neurons’ preferred spike phase. This preferred spike-phase might thus be important to encode information shared among areas of the network (Fries, 2015; Womelsdorf et al., 2014b). We tested this hypothesis by quantifying how much *Outcome, Prediction Error*, and *Outcome History* information is available to neurons at different phase bins relative to their mean phase. If the phase-of-firing conveys information, then differences in spike counts between conditions should vary systematically across phases, as opposed to a pure firing rate code that predicts equal information for spike counts across phase bins (**Figure 4A**) (Hawellek et al., 2016; Womelsdorf et al., 2012). **Figure 4B** shows an example neuron exhibiting phase-of-firing coding (with spikes from ACC and beta phases from STR). This neuron exhibits large firing increases on error trials compared to correct trials, but only when considering spikes near its preferred spike phase; on the other hand, firing at the opposite phase showed a weaker difference. To quantify the phase-of-firing code for all three information types, we selected for each neuron the frequency within 10-25 Hz that showed maximal spike-LFP synchrony, subtracted the mean (preferred) spike phase to allow for comparison between neurons, and binned spikes on the basis of the LFP beta phases. To prevent an influence of overall firing rate changes between phase bins, we adjusted the width of each of the six phase bins to have (on average) an equal spike count across bins. We then fitted a GLM to the firing rates of each phase bin separately to quantify the *Outcome, Prediction Error*, and *Outcome History* encoding for each phase bin, and compared this phase specific encoding to a null distribution after randomly shuffling the spike phases (*see* **Methods**). **Figure 4C** illustrates example neurons for which the encoding systematically varied as a function of phase (*see also* **Figure S4**). The example spike-LFP pair from **Figure 4B** encoded the trial *Outcome* significantly stronger than a rate code in spikes near [-π/2, π/2] radians relative to its preferred spike phase and weaker than a firing rate code at opposite phases (**Figure 4C**, *left*); Similar phase-of-firing encoding was evident for *Prediction Error* and *Outcome History* as independent variable (**Figure 4C**, *middle and right panels)*.

**Figure 4.**
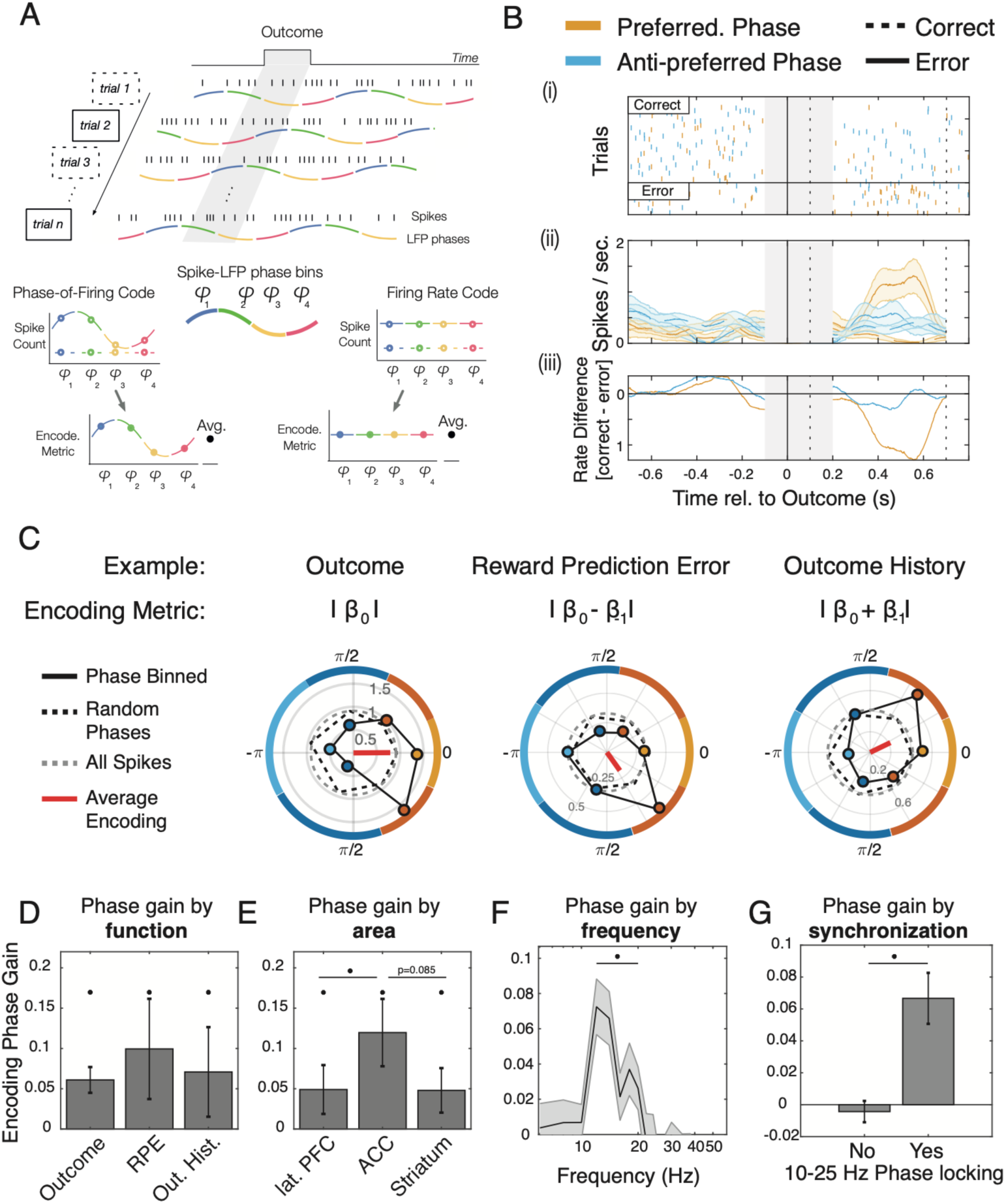
Encoded learning signals are modulated by beta phase. **(A)** Sketch of phase-dependent encoding analysis. Spikes were segregated according to the phase of the LFP, with bin limits adjusted such that an approximately equal number of spikes fell within each bin. Afterwards, a reduced GLM was run to extract the appropriate functional encoding metric for each phase bin. **(B)** Example cell showing phase-dependent outcome encoding. (i) Raster plot depicting the firing of the cell on correct (top) or error (bottom) trials. Color represents firing on the maximal (orange) or minimal (yellow) encoding phase. The grey box between [-0.1 0.2] represents spikes that were not analyzed due to this window not being used for phase estimation. (ii) Spike density function for correct/error trials, using spikes from the maximal/minimal encoding phase. (iii) The difference in firing rate between outcomes for the preferred and anti-preferred phase is visualized as a difference. The difference is greater on preferred rather than anti-preferred phases for an extended period of time. **(C)** Example of phase-of-firing encoding for Outcome, RPE, and Outcome History cells. Colored dots reflect the encoding metric for a corresponding phase bin. Zero radians corresponds to the preferred (mean) firing phase. Numbers on concentric circles are the value of the encoding metric. The grey dotted line represents the encoding metric estimated using all spikes, whereas the black dotted line represents the average across many permutations of spike phases. The red line is the average direction. Colored border lines represent size of the bin. These cells show stronger encoding near the 0 rad phase, and weaker encoding on opposite phases. **(D)** Median and standard error of the *EPG (Encoding Phase Gain)* in each functional cluster. All clusters showed some evidence of significant phase gain (Wilcoxon signrank test, one-sided, p<0.05). **(E)** Median phase gain for each area. ACC had a higher phase gain than LPFC or STR. **(F)** Median phase gain per frequency for encoding and locking cells (n=877). Phase gain was above chance at [10 20] Hz. **(G**) Median phase gain for spike-LFP pairs (with significant encoding) that showed significant synchrony in the 10-25 Hz beta band (n=877), vs. those that did not (n=2500). Non-locking cell showed a sign. less phase gain compared to locking cells (Wilcoxon ranksum test, p<0.05).

We estimated the strength of phase modulation of rate encoding for each spike-LFP pair as the amplitude of a cosine fit to the phase-binned encoding metric and normalized it by the cosine amplitude obtained from fitting the phase-binned metric after randomly shuffling spike phases. We refer to this difference of the observed to the randomly shuffled phase modulation of encoding as the *Encoding Phase Gain* (*EPG*). Of the 877 spike-LFP pairs that significantly synchronized in the 10-25 Hz and encoded information in their firing rate, we found that 139 (16%) spike-LFP pairs showed significant phase-of-firing encoding, i.e. these pairs encoded significantly more information in the phase of firing than in their firing rate alone (randomization test, p<0.05). A significant *EPG* was evident for neurons whose average firing encoded *Outcome* (Wilcoxon signrank test, p∼0), *Prediction Error*, (p=0.025), and *Outcome History* (p=0.016). The *EPG* did not differ between these three functional clusters (Kruskal Wallis test, p=0.86) (**Figure 4D**). Similarly, *EPG* was evident for spike-LFP pairs with the spiking neuron in ACC (p∼0), in STR (p=0.00028), and in LPFC (p=0.015) (**Figure 4E**). EPG differed between areas (Kruskal-Wallis test, p=0.016), with ACC showing significantly larger *EPG* than LPFC (p=0.004), and a similar trend when compared to STR (p=0.086). *EPG* differences were more pronounced when selecting for each encoding metric the 25% of spike-LFP pairs with the largest *EPG*. This selection revealed stronger EPG encoding of *Prediction Error* compared to *Outcome* (p∼0) and *Outcome History* (p=∼0). Large *EPG* were found significantly more likely in ACC than in LPFC (p=0.00066) or STR (p=0.032). These results were similar in each monkey (**Figure S3C**). Encoding designation was largely stable across phase bins, with ∼90% of spike-LFP pairs exhibiting similar beta coefficient signs across all phase bins. *EPG* did also not depend on the number of bins used (Spearman rank correlation, R=0.023, p=0.18). Likewise, *EPG* was similar for neurons that showed narrow (N) and broad (B) action potential waveforms that corresponds to putative distinct cell types with their encoding phase gain statistically indistinguishable in the ACC (N_N_=70, mean=-0.026 ± 0.029; N_B_=48, mean = -0.0047±0.06; Kruskal Wallis test for equal median, p=0.49), LPFC (N_N_=85, mean=0.11 ± 0.04; N_B_=54, mean = 0.0057±0.080; p=0.28), or STR (N_N_=37, mean=0.014 ± 0.08; N_B_=41, mean = 0.017±0.11; p=0.40) (see **Methods**).

### Phase-of-Firing Encoding Depends on Significant Synchronization in the Beta Band

We next tested how *EPG* related to the strength of synchronization. First, we found that *EPG* was significant at the population level and strongest in the same beta frequency band that showed the strongest spike-LFP synchronization (**Figure 4F**; Wilcoxon signrank test, p<0.05). Overall, *EPG* was most prevalent and significantly larger in spike-LFP pairs showing significant phase synchronization (**Figure 4G**; Kruskal-Wallis test, p∼0). These results indicate that *EPG* was evident when neurons encoded *Outcome, Prediction Error* and *Outcome History* in their firing rate *and* when they synchronized at beta band frequencies.

We next asked whether the synchronizing phase that carried information was endogenously generated or whether it was externally triggered by the reward onset. We calculated the *EPG* with and without subtracting the reward-onset aligned evoked LFP response (*see* **Methods**). We found that the *EPG* was not different with (median=0.0704 ± 0.012 SE) versus without (median=0.067 ± 0.016 SE) subtraction of the time-locked, evoked potential, suggesting that the beta oscillation events providing informative phases were endogenously generated (Kruskal-Wallis test, p=0.90). We also tested whether LFP power variations influenced the phase-of-firing encoding but found that the *EPG* did not correlate with beta band power variations (Spearman rank correlation, R=0.051, p=0.13).

### Preferred Spike Phase and Encoding Phase Differ for Prediction Error

The phase-of-firing encoding so far might be due to stronger spike-LFP synchronization in one task condition than in another condition (e.g. in error trials versus correct trials). Such site-specific selectivity of neuronal synchronization has been reported in previous studies (e.g. (Antzoulatos and Miller, 2016; Salazar et al., 2012; Womelsdorf et al., 2010). To test this possibility, we compared spike-LFP synchronization (indexed with the PPC), in those two trial conditions that were predicted to have the maximal firing rate difference. For *Outcome* encoding we compared correct vs error trials; for *Prediction Error* encoding we compared correct trials following error *versus* error trials following correct trials; and for *Outcome History* encoding we compared correct trials following correct, or errors following errors. We then correlated the absolute difference in PPC in the beta band between two conditions with the phase-of-firing encoding. We found statistically indistinguishable levels of PPC between conditions for neurons encoding *Outcome* (Spearman rank correlation, R=0.038, p=0.35), *Prediction Error* (R=0.050, p=0.58), or *Outcome History* (R=-0.027, p=0.76).

This result suggests that the strength of phase-of-firing encoding does not follow simply from differentially strong spike-LFP phase synchronization. Rather, the spike-phase at which neurons maximally synchronize might not always coincide with the spike-phase at which encoding of task variables is maximal. Indeed, we often observed that the phase with maximal encoding was not at the zero-phase bin, i.e. it deviated from the preferred spike-phase (**Figure 4C; Fig S4)**. We tested this scenario by first calculating the preferred spike-phase for each neuron, and then quantifying the phase with maximal encoding relative to that phase. We found that all encoding neurons synchronized on average at similar phases, above what would be expected by chance (Hodges-Ajne test; *Outcome*, average phase: -0.28 ± 0.0034 SE radians, p∼0; *RPE*. average phase: 0.35 ± 0.0034 SE radians, p=0.00084; *Outcome History*. average phase: -0.68 ± 0.0045 SE radians, p=0.0013) (**Figure 5A**). The preferred spike-phase differed between the three encoding classes (Watson Williams test, p∼0; each pairwise comparison showed: Watson-Williams test, p<0.02; **Figure 5B**).

**Figure 5.**
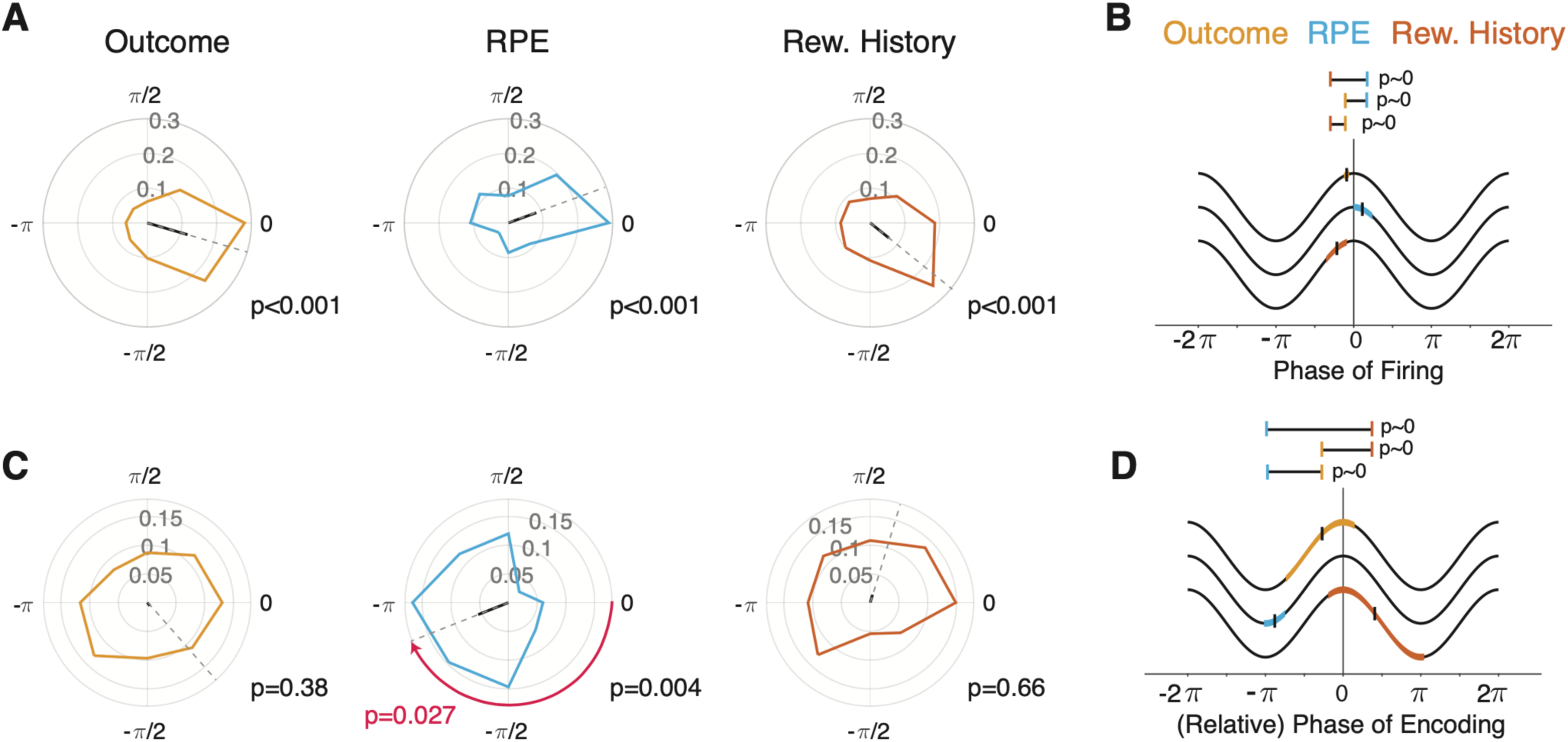
Preferred firing and encoding phase are dissociated in RPE cells. **(A)** Proportion of cells with a preferred phase of firing for each encoding cluster. Black lines depict the average phase. *Outcome (yellow), RPE (blue)* and *Outcome History* (*red*) cells showed strong evidence of phase concentration (Hodjes-Ajne test, p<0.05). **(B)** Mean preferred phase of firing for *Outcome, RPE*, and *Outcome History* clusters. Colored patches represent the 95% confidence interval about the mean, depicted as a black horizontal line. The distribution of preferred firing phases differed between all encoding clusters (Watson-Williams test, p<0.05). **(C)** Relative encoding phase (where 0 corresponds to the preferred phase). *Outcome* and *Outcome History* cells did not show an encoding phase preference, whereas RPE cells did (Hodges-Ajne test, p=0.0004). Importantly, for these RPE cells, this phase differed significantly from 0 (the preferred phase of firing; Median test, p=0.027) **(D)** Similar conventions as in (B) but for the encoding phase. The distribution of encoding phases differed between all three clusters.

Next we quantified for each cluster whether the phases showing maximal encoding were consistent across spike-LFP pairs (**Figure 5C**). To this end, we extracted the phase offset from our cosine fit, which represents the phase at which encoding was maximal relative to the preferred spike-phase. *Outcome* encoding neurons showed a preferred encoding phase that randomly varied across neurons (Hodges-Ajne test, p=0.38), as did *Outcome History* neurons (p=0.66) (**Figure 5C**). In contrast, *Prediction Error* encoding neurons significantly encoded at similar phase-offsets relative to the neuron’s synchronizing phases (Hodges-Ajne test, p=0.0004 at -2.76 ± 0.047 SE radians, corresponding to 27 ms at a 15 Hz oscillation cycle), which was significantly different than the mean spike phase (Median test, p=0.027). This effect was particularly pronounced for *RPE* cells in ACC **(Figure S5)**, and was consistent across both monkeys (**Figure S3D)**. The phases showing maximal encoding differed between all three functional clusters (Watson-Williams test; p<0.001; **Figure 5D**). These result show that the preferred spike phase and the encoding phases are dissociated from one another for all types of information. Additionally, for *Prediction Error* information there was a systematic offset of maximal encoding at ∼27 ms away from the preferred (mean) spike phase. There was no relation between the phase of maximal encoding and the number of bins used (Circular-Linear correlation, R=0.013, p=0.76).

We next validated that the dissociation of spike- and encoding-phases was not influenced by phase shifts due to differences in the peak oscillation frequencies within the beta band. The three sets of neuronal encoding clusters synchronized on average at the same center frequency of ∼15 Hz (Kruskal Wallis, p=0.34), and showed maximal phase-of-firing encoding at similar frequencies (also ∼15Hz) (Kruskal Wallis test, p=0.23). Moreover, the frequency showing strongest spike-LFP synchronization and the frequency showing maximal encoding phase gain matched closely (median frequency ratio: 1± 0.019 SE). This similarity of synchronization and encoding frequency did not differ on the basis of the functional designation (Kruskal-Wallis test, p=0.17), nor the area from which the spikes were sampled (Kruskal Wallis test, p=0.55).

## Discussion

### Summary

Here, we found a substantial proportion of neurons whose phase-of-firing in the beta frequency band conveyed significantly more information about three learning variables than their firing rate alone. This encoding phase gain was evident for spikes generated within the ACC, LPFC and STR of nonhuman primates in a [0.1 0.7] second period of outcome processing during reversal learning performance. Phase-of-firing encoding was most prominent at the 10-25 Hz beta frequency at which spikes synchronized to the local fields across areas. However, the strength of spike-LFP phase synchronization could not explain the strength of the phase-of-firing encoding. Rather, maximal encoding occurred for many neurons at phases away from the preferred spiking phase. The dissociation of spiking- and encoding phases was particularly prominent for information about the *Prediction Error*.

Taken together, these results provide a first report of information multiplexing of learning variables at segregated phases of a beta oscillation that is synchronized across medial and lateral fronto-striatal loops. These findings suggest that oscillation phases are important carriers of information, above and beyond that of a phase-blind firing code. The gain of information through phase-specific firing provides an intriguing dynamic code that could link principles of efficient neuronal information transmission with the demands of representing multiple types of information in the same dynamical neural system.

### Distributed Encoding of Learning Variables at a Shared Beta Rhythm Frequency

We found that three critical variables needed for adjusting behavior are represented in segregated neuronal populations not only in their firing rates, but in phase specific firing at a beta frequency that is shared among ACC, LPFC and STR. This finding suggests that the beta frequency could serve as a primary carrier for the fast distribution of learning related information within fronto-striatal networks (Spitzer and Haegens, 2017; Womelsdorf and Everling, 2015). Prior studies have shown that ACC, LPFC and STR causally contribute to fast learning of object values. With lesioned ACC, rhesus monkeys fail to use outcome history for updating values and show perseverative behaviors (Kennerley et al., 2006). Without LPFC, rhesus monkeys fail to recognize when a previously irrelevant object becomes relevant, as if they fail to calculate prediction errors needed for updating their attentional set (Rudebeck et al., 2017). When the anterior STR is lesioned, nonhuman primates tend to stick to previously learned behavior and show a lack of sensitivity to reward outcomes (Hikosaka et al., 2017, 2019). These behavioral lesion effects are consistent with the important role of each of these brain areas to track the history of recent outcomes, registering newly encountered (current) outcomes, and calculating the unexpectedness of experienced outcomes (prediction error). Consequently, our finding of segregated neuronal ensembles encoding *Outcome, Prediction Error* and *Outcome History* complement a large literature that documents how these variables are represented in the firing of neurons across fronto-striatal areas.

What has been left unanswered, however, is how this firing rate information about multiple variables emerges at similar times and similar proportions across areas. Prior studies suggest that firing rate correlations between brain areas are relatively weak and poor candidates for veridical information transfer (Hahn et al., 2019; Kumar et al., 2010; Oemisch et al., 2019), while temporally aligning the spike output of many neurons to the phases of precisely timed, synchronized packets are a theoretically, particularly powerful means in affecting postsynaptic neuronal populations (Azouz and Gray, 2003; Fries, 2015; Hahn et al., 2019; Luczak et al., 2015; Voloh and Womelsdorf, 2016). Our findings support this notion of a temporal code using synchronized oscillations by showing that those neurons that carry critical information in their firing rates also tend to synchronize long-range between ACC, LPFC, and STR at a shared 10-25 Hz beta frequency. This beta frequency is thus a powerful candidate for distributed information transfer, because spike output of many neurons is concentrated at the same phase and thus activate postsynaptic membranes at similar times. This scenario of beta rhythmic information exchange within fronto-striatal networks is supported by previous nonhuman primate studies that demonstrated 10-25 Hz beta rhythmic synchronization during active task processing states between ACC and LFPC (Smith et al., 2019; Voloh and Womelsdorf, 2017a; Womelsdorf et al., 2014a), between PFC and STR (Antzoulatos and Miller, 2014), between ACC and FEF (Babapoor-Farrokhran et al., 2017), between LPFC and FEF (Antzoulatos and Miller, 2016; Salazar et al., 2012), and between lateral PFC or FEF with posterior parietal cortex (Antzoulatos and Miller, 2016; Buschman and Miller, 2007; Buschman et al., 2012; Dean et al., 2012; Salazar et al., 2012). Each of these studies have shown short-lived rhythmic long-range synchronization between distant brain areas during cognitive tasks at a similar ∼15 Hz frequency. Our findings critically complement these studies by revealing that 10-25 Hz spike-LFP synchronization is prevalent not only during cognitive processing, but also during the processing of outcomes after attention has been deployed and choices have been made. During this post-choice outcome processing, fronto-striatal circuits are likely to adjust their synaptic connection strength to minimize future prediction errors and improve performance (Hikosaka et al., 2019; Leong et al., 2017; Oemisch et al., 2019). Our results suggest that this updating utilizes beta rhythmic activity fluctuations during the post-choice outcome processing period.

### Multiplexing of Information Through Phase-of-Firing Encoding

Our finding that spiking output carries separable types of information at different phases of the same oscillation frequency has potentially far-reaching implications. By finding that *Outcome, Prediction Error and Outcome History* were encoded at separate phases, the neuronal spiking activity effectively multiplexes independent information streams at different phases of beta synchronized firing. This stands in contrast to prior studies reporting that long-range beta rhythmic synchronization between LPFC, ACC or STR in the primate encoded relevant task variables via the strength of beta synchrony (Antzoulatos and Miller, 2016; Babapoor-Farrokhran et al., 2017; Buschman and Miller, 2007; Buschman et al., 2012; Dean et al., 2012; Salazar et al., 2012; Womelsdorf et al., 2014a). For example, some prefrontal cortex neurons synchronize stronger at beta to posterior parietal areas when subjects choose one visual category over another (Antzoulatos and Miller, 2016), or when they maintain one object over another in working memory (Salazar et al., 2012). These findings are consistent with a communication-through-coherence schema where upstream senders are more coherent with downstream readers when they successfully compete for representation (Bosman et al., 2012; Fries, 2015; Womelsdorf et al., 2007). Yet it has remained unclear how such a scheme may operate when multiple items must be multiplexed and transmitted in the same recurrent network (Akam and Kullmann, 2014; Khamechian et al., 2019; Kopell et al., 2011; Palminteri et al., 2015; Siegel et al., 2009). Computationally, the multiplexing and the efficient transmission of information can operate in tandem when the temporal organization of activity is exploited at the sending and receiving site (Buzsáki, 2010; Hahn et al., 2019; Kumar et al., 2010; Luczak et al., 2015). Consequently, selective synchronization between distal sites could be leveraged to enhance transmission *selectivity*, whereas temporally segregated information streams could enhance transmission *capacity* (McLelland and VanRullen, 2016). Our results resonate with this view by showing that neurons that synchronize long-range at one oscillation phase carries information of any of three learning variables at phases systematically offset from the synchronizing phase.

By finding evidence for such a temporal multiplexing in the beta frequency band we critically extend previous reports of phase encoding of information for object features, object identities, and object categories at theta, alpha and gamma frequencies (Kayser et al., 2012; Siegel et al., 2009; Turesson et al., 2012; Womelsdorf et al., 2012). In our study, the beta rhythmic phase-of-firing multiplexing applied to complex learning variables that were needed to succeed in the behavioral learning task. In particular, the presence of reward prediction error information provides a critical teaching signal that indicates how much synaptic connections should change to represent future value expectations more accurately (Leong et al., 2017; Oemisch et al., 2019). Our results suggest that this updating can utilize spike-timing dependent plasticity mechanisms that are tuned to firing phases ∼27 ms away from the preferred synchronization phase in the beta frequency band. How such a temporal organization in the beta band is used in the larger fronto-striatal network will be an important question for future studies.

So far, evidence in humans and rodents suggested that processes linked to beta frequency activity during the evaluation of outcomes support the detection of errors and the updating of erroneous internal predictions (Howe et al., 2011; Tan et al., 2014b). In fact, there have been conflicting views on whether beta oscillations related to outcome signals are more likely to reflect a weighted integration of recent outcomes, or the unexpectedness of the current, observed outcome relative to recent outcomes (Howe et al., 2011; Tan et al., 2014a). Our findings reconcile these viewpoints by documenting that encoding of *Outcome History* weights and of *Prediction Errors* coexist in the same circuit at the same oscillation frequency in phase dependent firing of single neurons.

### Neuronal Mechanisms underlying Phase-of-Firing Multiplexing

We found that the beta phase allowing maximal encoding of *Prediction Errors* was offset ∼27 ms on average from the phase at which most spikes synchronized to the local fields. Such a dissociation of spike-phase and encoding-phase has been reported previously for the beta frequency band in parietal cortex, where maximal information of joint saccadic and joystick choice directions were best predicted by spike counts at ∼50 degree away from the preferred beta spike phase (Hawellek et al., 2016). Such phase offsets underlying maximal encoding in parietal cortex as well as in ACC, LFC and STR in our study provide constraints on the possible circuit mechanisms that permit temporal segregation of inputs streams through phase specific oscillatory dynamics (Akam and Kullmann, 2014). One possible circuit mechanism that implements and utilizes multiplexed information streams through phase specific firing has been described and computationally modelled specifically for the low 10-20 Hz frequency range (Gelastopoulos et al., 2019; Kopell et al., 2011). This work suggests that distinct sets of pyramidal neurons can encode distinct input streams in their firing phases at 10-20 Hz beta phases when these inputs streams arrive with a phase offset to each other, e.g. when they arrive sequentially in time. According to this schema, a first input stream activates pyramidal neurons in deep cortical layers that feed information to superficial layers whose interlaminar inhibitory connections closes an interlaminar reverberant loop of activity. This interlaminar ensemble follows a beta activity rhythm due to cell specific dynamics that maintains the beta-phasic firing of active neurons (Kopell et al., 2011; Womelsdorf et al., 2014b). When a second input stream activates another set of pyramidal cells within the same beta rhythmic neural population, the input timing of that second stream was maintained at a different phase than the phase of the first activated ensemble (Gelastopoulos et al., 2019). The parallel coding of information at a common beta rhythm in these models provide a qualitative proof of concept about phase specific encoding of multiple types of inputs in larger beta rhythmic ensembles, and suggests their mechanistic realization (Gelastopoulos et al., 2019). Based on our results we predict that the set of neurons encoding current *Outcomes, Prediction Errors* and *Outcome History* will emerge earliest in deep ACC and LPFC layers that are associated with prominent beta rhythms (Bastos et al., 2018) and then transmit to superficial layer neurons to form an interlaminar beta rhythmic ensemble multiplexing different types of information at the same shared beta frequency oscillation. Future work needs to specify how such multiplexing ensembles can transform inputs to generate novel representations.

In summary, we have documented that learning variables are encoded at separable phases of firing of neurons that synchronize long–range across primate fronto-striatal circuits. These phase encoding neurons also carried information in overall firing rate modulations which clarifies that an *asynchronous* rate code and a *synchronous* temporal code coexist in the same circuit (Kumar et al., 2010). By exploiting the temporal structure endowed in long–range neuronal synchronization our findings suggest how neuronal assemblies in one brain area could be read out from neural assemblies in distally connected brain areas (Singer and Gray, 1995). This phase-of-firing schema entails key features required from a versatile neural code including the efficient neural transmission and the effective representation of variables needed for adaptive goal directed behavior (Perkel and Bullock, 1968).

## Acknowledgments

This research was supported by a grant from the Canadian Institutes of Health Research (T.W.) CIHR Grant MOP_102482 and by the National Institute Of Biomedical Imaging And Bioengineering of the National Institutes of Health under Award Number R01EB028161 (T.W.). The content is solely the responsibility of the authors and does not necessarily represent the official views of the Canadian Institutes of Health Research or the National Institutes of Health. We thank Dr. Andrew Tomarken for very useful methodological discussions.

## Author Contributions

Conceptualization, B.V. and T.W.; Methodology, B.V. and T.W.; Software, B.V. and T.W; Investigation, M.O. and T.W.; Writing – Original Draft, B.V. and T.W..; Writing – Review & Editing, all authors; Visualization, B.V. and T.W..; Supervision, T.W.

## Declaration of Interests

The authors declare no competing interests.

## STAR Methods

### Animals

Data was collected from two adult, 9 and 7-year-old, male rhesus monkeys (*Macaca mulatta*) following procedures described in (Oemisch et al., 2019). All animal care and experimental protocols were approved by the York University Council on Animal Care and were in accordance with the Canadian Council on Animal Care guidelines.

### Behavioral paradigm

Monkeys performed a feature-based reversal learning task that required covert attention to one of two stimuli based on the reward associated with the color of the stimuli. Which stimulus color was rewarded remained identical for ≥30 trials and reversed without explicit cue. The reward reversal required monkeys to utilize trial outcomes to adjust to the new color-reward rule. Details of the task have been described before (Oemisch et al., 2019). Each trial started when subjects foveated a central cue. After 0.5-0.9 sec, two black and white gratings appeared. After another 0.4 sec., the stimuli either began to move within their aperture in opposite directions (up-/downwards) or were colored with opposite colors (red/green or blue/yellow). After another 0.5-0.9 sec, they gained the color when the first feature was motion, or they gained motion when the first feature had been color. After 0.4-0.1 sec, the stimuli could transiently dim. The dimming occurred either in both stimuli simultaneously, or separated in time by 0.55 sec. Dimming represented the go-cue to make a saccade in the direction of the motion when it occurred in the stimulus with the reward associated color. The dimming acted as a no-go cue when it occurred in the stimulus with the non-rewarded. A saccadic response was only rewarded when it was made in the direction of motion of the stimulus with the rewarded color. Motion direction and location of the individual colors were randomized within a block. Thus, the only feature predictive of reward within a block was color. Color-reward associations were constant for a minimum of 30 trials, a maximum of 100 trials, or until 90% performance was reached (running average, 12 trials). The block change was uncued. Rewards were deterministic.

### Electrophysiology

Extra-cellular recordings were made with 1–12 tungsten electrodes (impedance 1.2–2.2 MOhm, FHC, Bowdoinham, ME) in anterior cingulate cortex (ACC; area 24), prefrontal cortex (LPFC; area 46, 8, 8a), or anterior striatum (STR; caudate nucleus (CD), and ventral striatum (VS)) through a rectangular recording chambers (20 by 25 mm) implanted over the right hemisphere (**Figure S1**). Electrodes were lowered daily through guide tubes using software-controlled precision micro-drives (NAN Instruments Ltd., Israel and Neuronitek, Ontario, Canada). Data amplification, filtering, and acquisition were done with a multichannel acquisition system (Neuralynx). Spiking activity was obtained following a 300–8000 Hz passband filter and further amplification and digitization at 40 kHz sampling rate. Sorting and isolation of single unit activity was performed offline with Plexon Offline Sorter, based on analysis of the first two principal components of the spike waveforms. Experiments were performed in a custom-made sound attenuating isolation chamber. Monkeys sat in a custom-made primate chair viewing visual stimuli on a computer monitor running with a 60 Hz refresh rate. Eye positions were monitored using a video-based eye-tracking system (EyeLink, SRS Systems) calibrated prior to each experiment to a nine-point fixation pattern. Eye fixation was controlled within a 1.4°–2.0° radius window. During the experiments, stimulus presentation, monitored eye positions, and reward delivery were controlled via MonkeyLogic (www.brown.edu/Research/monkeylogic/). Liquid reward was delivered by a custom-made, air-compression controlled, and mechanical valve system. Recording locations were aligned and plotted onto representative atlas slices (Calabrese et al., 2015).

### Data Analysis

Analysis was performed with custom Matlab code (Matlab 2019a), using functions from the fieldtrip toolbox (http://www.fieldtriptoolbox.org). For Elastic-net regression, the *glmnet* package in R was used (Friedman et al., 2010). Only correct and error responses were analyzed. Error responses included those where the responses were made to the incorrect target, or in the incorrect response window. Only trials from learned blocks were included, with a minimum of two blocks, unless otherwise indicated. Learned blocks were defined as ones where animals reached 90% correct responses within the last 10 trials within the block. Standard errors of the median were estimated via bootstrapping (200 repetitions, unless otherwise indicated)

### Behavioral analysis

To determine the timescale over which past outcomes are integrated, we used a binomial GLM:

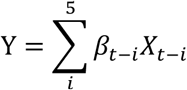

Where Y was the current outcome, B_t-i_ is the influence of outcome X_t-i_ on trial t-i. Outcome for trial t-5 was defined as a nuisance variable that accounted for all responses occurring over very long time-scales (similar to (Rudebeck et al., 2017; Walton et al., 2010)).

### Rate encoding of outcome history

To test how individual units integrated outcome history, we used a Poisson GLM:

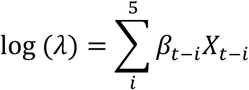

Where λ was the conditional intensity (spike count), B_t-i_ is the influence of outcome X_t-i_ on trial t-i. Firing counts on each trial were determined in a [0.1 0.7]s window after outcome onset (Oemisch et al., 2019). Neurons were included in the analysis if they were isolated for more than 25 (learned) trials across at least two blocks, and if they showed an overall firing rate of >1 Hz.

To mitigate issues of multi-collinearity, and extract only the most predictive regressors, we employed penalized LASSO regression, which shrinks small coefficients to zero, and tends to select one regressor in the presence of collinearity (Friedman et al., 2010; Tibshirani, 1996). We used the R package *glmnet*, with an alpha of 0.95. Optimal lambdas were the minimum as selected by 10-fold cross validation. To assess model stability and extract significant fits, we used a bootstrap approach, whereby trials were sampled with replacement 1000 times and the procedure was rerun. As the LASSO shrinks non-valuable predictors to zero, a model fit was said to be significant if at least one regressor (outcome t-4 to t-0) was non-zero more than 95% of the time.

### Functional clustering based on neural encoding

To determine the putative function of significantly encoding units, we used a clustering approach via bootstrapped K-means. We clustered cells on the basis of their mean beta weights as determined by the penalized regression model (see above). As a preprocessing step, for units where the current outcome was negatively encoded (i.e. encodes errors), we flipped the sign of every coefficient in that model. This has the effect of erasing the directionality of any functional association, and thus collapses neurons with similar functions (for example, *Error* or *Correct* encoding units become *Outcome* encoding units). Cells were independently clustered for each area.

We clustered cells using K-mean clustering with a cosine distance metric (which is insensitive to the magnitude of the vector and is instead concerned with the direction). To determine the optimal number of clusters, we used the Silhouette method with bootstrapping. Briefly, cells were sampled with replacement 1000 times and for each iteration, the optimal number of clusters was extracted where the silhouette was maximal. The overall optimal number of clusters *Nc* was the mode over all bootstrap iterations.

We clustered cells on the basis of their clustering stability. To do so, we built a similarity matrix via a bootstrap approach. First, we resampled with replacement individual cells. Next, we ran K-means with cosine distance and *Nc* clusters. For units that were clustered together, their respective cell in the similarity matrix was incremented by one. Because bootstrapping could sample the same units twice, these pairs were ignored. Bootstrapping was run 100000 times. The similarity matrix S was defined as the proportion of times that pairs of cells were clustered together. We then formed a dissimilarity matrix D=1-S, and computed the final cluster assignment via agglomerative clustering with Euclidian distance and *Nc* clusters.

### Metrics for outcome, outcome history, and prediction error

We quantified the degree of encoding of *Outcome* (E_outcome_), *Outcome History* (E_history_), and *Prediction Error* (E_pe_) on the basis of the GLM weights for trials -1 and 0:

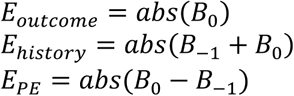

We refer to these generically as *Encoding Metrics*.

### Latency analysis

To determine the latency of encoding for each functional cluster, we performed a time-resolved analysis (**Figure S1C,D**). On the basis of our previous results showing that the outcome on trial 0 and -1 were most predictive (**Figure 2**), we used a simpler GLM of just the current and previous outcome. For the response variable, we calculated the spike density using a sliding Gaussian window, with a 200 ms window and 50 ms standard deviation. We performed this analysis [-0.4 0.7] around outcome onset. We thus obtained a time-resolved estimate of encoding.

To determine the latency of significant encoding, we looked at time points in the post-outcome period that were significantly above the pre-outcome period. First, we z-score normalized each individual encoding metric to the pre-outcome period ([-0.4 0] sec). Next, we asked, for each time point, whether the population response was significantly different from zero via a Wilcoxon signrank test. We then extracted the largest cluster mass of contiguous significant time points (e.g. (Maris and Oostenveld, 2007)) to find a *time-of-interest*. We then extracted, for each individual cell, the time point where the cumulative encoding metric reached a 10% threshold within this time-of-interest. Thus, we obtain for each encoding cluster a distribution of latencies of when they started to show significant encoding of *Outcome, Outcome History* or *Prediction Error*.

### Spectral decomposition and spike-LFP phase synchronization

We focused our analysis on distal recording sites, thus obviating any concerns of spike energy bleeding into the LFP (Zanos et al., 2011). We considered frequencies from 6 Hz to 60 Hz. For frequencies from 6-30Hz, the resolution was 1 Hz, and above that it was 2Hz. For every frequency F, we determined the spike-LFP phase by extracting an LFP segment centered on the spike of length 5/F (i.e. 5 cycles). Spectral decomposition was done via an FFT after applying a Hanning taper. This procedure was applied separately to the pre-outcome period [-1 0]sec, and the post-outcome period [0.1 1] sec.

The strength of spike-LFP synchronization was quantified using the pairwise phase consistency (PPC), which is unbiased by the number of spikes (Vinck et al., 2011). The PPC is quantified on the basis of pairwise differences between spike-phases. If spikes tend to fire on specific phases, phase difference will be concentrated, and thus the PPC will take on a high value, whereas if spikes are distributed randomly relative to the LFP phase, phase differences will be random and the PPC will tend towards zero.

The effect size was determined as previously reported (Voloh and Womelsdorf, 2017b; Womelsdorf et al., 2014c):

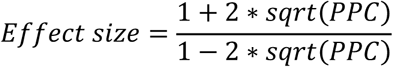

This effect size can be interpreted as the relative increase in spike rate at the cell’s preferred (mean) phase over its anti-preferred (opposite) firing phase. For example, a PPC value of 0.01 corresponds to a 1.5 times greater spike rate at the preferred phase.

We determined the frequency at which spike-LFP phase synchronization was significant by determining peaks in the PPC spectrum. A cell was said to synchronize to a particular frequency if the following criteria were met: (1) Peaks had to be above a threshold of 0.005, (2) show a minimum prominence of 0.005, and (3) show significant Rayleigh test (i.e. phase concentration).

To test for inter-areal differences in spike-beta synchronization, we extracted the PPC at the frequency in the [10 25] Hz band that showed significant encoding. For those cells that did not show such encoding, we extracted the frequency of the maximal PPC in this band instead. We tested for differences in synchronization strength using a one-way ANOVA, and report on pairwise comparisons after multiple comparison correction.

### Phase-of-Firing dependent encoding of Outcome, Outcome History and Prediction Error

To determine if spikes falling on certain phases of the LFP were more informative, we re-ran the (reduced) GLM on phase-binned spikes, using only the previous and current outcomes (see latency analysis). We first aligned all spike-triggered-phases to the circular mean of their distribution. Phases were extracted from the frequency of the corresponding maximal peak in the [10 25] Hz band in the PPC. However, if spikes are phase locked to an LFP, the firing rate around the preferred phase will naturally be higher. Thus, we used non-equal bin sizes, adjusting the bin limits such that they contained approximately equal numbers of spikes, and phase-labelled individual spikes accordingly. We then re-ran the GLM analysis on spikes falling within a particular bin and computed the encoding metrics as described previously. To aid in comparison, we also fit the model using randomly permuted phases (thus preserving the over-all rate response structure). We ignored spike-LFP pairs where the GLM could not converge to a solution and threw a warning, or where the beta coefficients were above 20 (however, relaxing or tightening this constraint did not change the results).

To determine if the phase and degree of phase-dependent encoding, we fit a cosine function to the phase-binned encoding values (illustrated in **Figure 4A**) (Siegel et al., 2009; Womelsdorf et al., 2012). From this fit, we obtain three values: T (phase offset, or phase of cosine maxima), A (amplitude), and M (overall mean, or offset). The value T is thus the phase at which encoding is maximal, relative to the preferred firing phase. To compare the strength of encoding across functional clusters, we computed the *empirical phase-of-firing gain*:

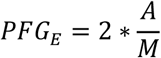

This quantity represents the difference in encoding between the peak and trough relative to the overall encoding strength. A PFG value of 0 implies that phase-of-firing adds no information (corresponding to a pure rate code), whereas PFG=1 means that encoding between the peak and trough is 100% stronger compared to the overall encoding strength. To determine if phase significantly added information above that of a rate code, we applied randomization. For each cell, we first permuted the phase label of each spike, re-ran the GLM, re-fit a cosine and extracted the phase-of-firing gain. This procedure was repeated 50 times. The randomized phase-of-firing gain PFG_r_ was the mean of this distribution. We report on the “excess” phase-of-firing gain, defined as the difference between empirical and randomized phase-of-firing gain, which we refer to in the manuscript at the *Encoding Phase Gain (EPG):*

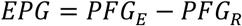

A positive value implies that encoding is modulated by phase above what would expected by chance.

We tested the stability of encoding across phase bins for each neuron (with significant rate encoding) by determining the sign of the encoding metric. We found that for the vast majority of cell-LFP pairs (∼90%), the sign of the encoding metric (ie before taking the absolute value) was the same for all 6/6 phase bins as for the full model.

To test the frequency specificity of the *EPG*, we extended the above analysis to the larger 6-60 Hz frequency range (**Fig 4F**). We statistically tested the *EPG* across frequencies using the Wilcoxon signrank test. We also tested the degree to which our results may be influenced by cue-aligned activity. To this end, we first obtained the average evoked potential for each LFP channel, and subtracted this component from individual trials. We then performed all steps of the analysis again to compare the original *EPG* with the *EPG* free from potential cue-aligned biases.

To test whether the preferred firing phase or relative phase with maximal encoding was concentrated above what would be expected by chance, we used the circular Hodges-Ajne test **(Figure 5)**. To determine whether the the phase showing maximal encoding differed from the preferred firing phase in each functional encoding cluster, we performed the Median test to test if the phase differed from zero (Zar, 2010) (**Fig. 5B**).

We tested how the strength of phase synchronization related to the strength of phase-of-firing encoding by performing two analysis. First, we compared encoding in cells that showed significant spike-phase synchronization to those that did not. For non-synchronizing cells, we selected the center frequency with the maximal PPC in the [10 25] Hz range, and computed the *EPG* at this frequency. We compared *EPG* between locking and non-locking populations using the Kruskal Wallis test (**Figure 4G)**. Second, we asked whether spike-phase synchrony in different trial conditions contained similar information to that of the phase-of-firing. To this end, for each encoding cell, we compared trials that were predicted to have the maximal firing rate differences. For *Outcome* encoding we compared correct *versus* error trials. For *Prediction Error* encoding we compared correct trials following error *versus* following error trials following correct. For *Outcome History* cells, this was errors followed by errors *versus* correct outcomes followed by correct. We took the absolute difference of the PPC between the two conditions, and correlated it with the *EPG* of the respective cell using the Spearman rank correlation.

We also tested whether phase gain depended on the number of bins used to fit the cosine function. We performed the analysis for 4, 6, 8, and 10 bins. We used Spearman rank correlation to determine if *EPG* was related to the number of bins, and circular-linear correlation to associate the phase of maximal encoding with the number of bins (Zar, 2010).

### Cell-type classification and analysis

To determine if phase-modulated encoding of information differed based on cell type, we focused the following analysis on highly isolated single units that showed encoding of learning-relevant variables and significant, prominent spike-beta locking. Detailed information is provided in (Oemisch et al., 2019). Briefly, to distinguish putative interneurons (narrow-spiking) and putative pyramidal cells (broad-spiking) in LPFC and ACC, we analyzed the peak-to-trough duration and the time for repolarization for each neuron. After applying Principal Component Analysis (PCA) using both measures, we used the first principal to discriminate between narrow and broad-spiking cells. This allowed for better discrimination than using either measure alone. We confirmed that a two-Gaussian model fit the data better than a one-Gaussian model using the Akaike and Bayesian Information Criterion (AIC, BIC). We then used the two-Gaussian model to define narrow and broad-spiking populations.

A similar analysis was applied to striatal units to distinguish putative interneurons from medium spiny projection neurons (MSN). Here, we use the peak-width and Initial Slope of Valley Decay (ISVD) (Berke, 2008; Lansink et al., 2010):

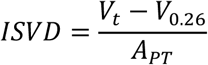

where V_T_ is the most negative value (trough) of the spike waveform, V_0.26_ is the voltage at 0.26 ms after V_T_, and A_PT_ is the peak-to-trough amplitude (Lansink et al., 2010). After PCA and two-Gaussian modelling (as described above), we defined two cut-off points. The first cutoff was the point at which the likelihood of narrow spiking cells was 3 times larger than the likelihood of broad-spiking cells, and vice-versa for the second cutoff.

We compared differences in *Encoding Phase Gain* between narrow and broad spiking neurons using the Kruskal-Wallis test, independently for each area. To clarify, we analyzed spike-LFP pairs here; thus, the same neuron may be included more than once.

## SUPPLEMENTAL FIGURES

**Supplemental Figure S1.**
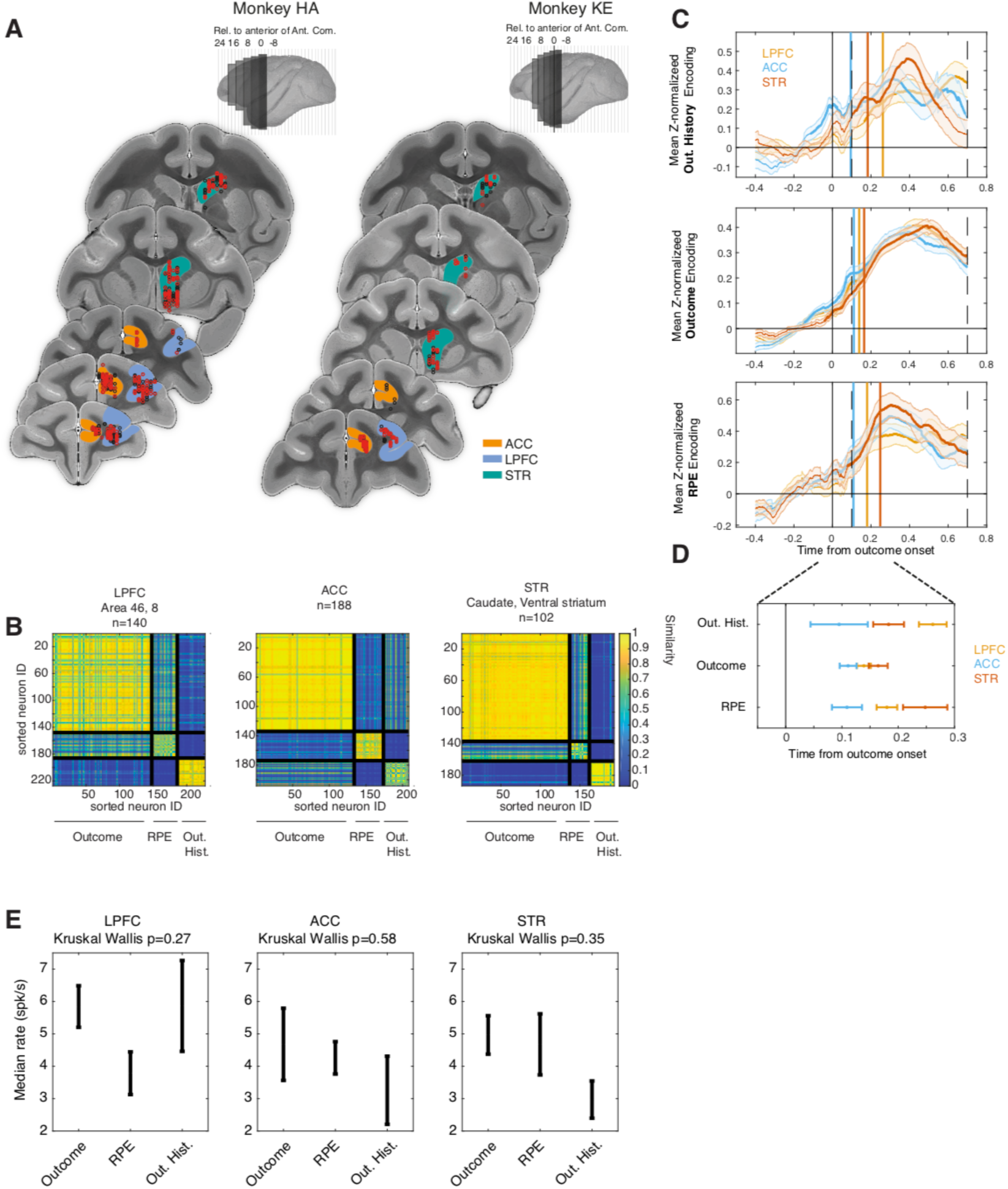
Recording sites and functional clustering. (**A**) All recording units used in the analysis. Units were collapsed across the anterior-posterior axis into equally spaced bins for each monkey. (*top*) Slice bin limits are visualized in the top, lateral view. (*bottom*) Units depicted on a representative atlas slice. Red dots represent encoding cells, and black represents non-encoding cells. Colored outlines correspond to the ACC (orange), STR (green), and LPFC (blue). Encoding cells were broadly distributed across the fronto-striatal axis. (**B**) Three encoding clusters emerge across the fronto-striatal axis. Pairwise similarity between pairs of neurons for each region. Neurons were sorted such that those in the same cluster were adjacent. (**C**) Time resolved, z-score normalized encoding metrics relative to the [-0.4 0] pre-outcome period, separated for *Outcome History* clusters (top), *Outcome* clusters (middle), and *RPE* clusters (bottom), and for LPFC (yellow), ACC (blue) and STR (red). At each time point, we assessed if encoding is above the baseline period (Wilcoxon signrank test). The bolded lines represent the largest contiguous mass where encoding was above baseline. This region represents the time-of-interest over which the latency was calculated for each individual cell. Latency was defined as the point at which 10% area-under-the-curve for the TOI was reached. Vertical lines depict the median latency for each cluster. **(D)** Median and standard deviation for each cluster of cells. All clusters showed significant encoding after the outcome onset. **(E)** Median and standard error of firing rate of encoding clusters for each of three regions. Fire rate differences were similar within all clusters (Kruskal

**Supplemental Figure S2.**
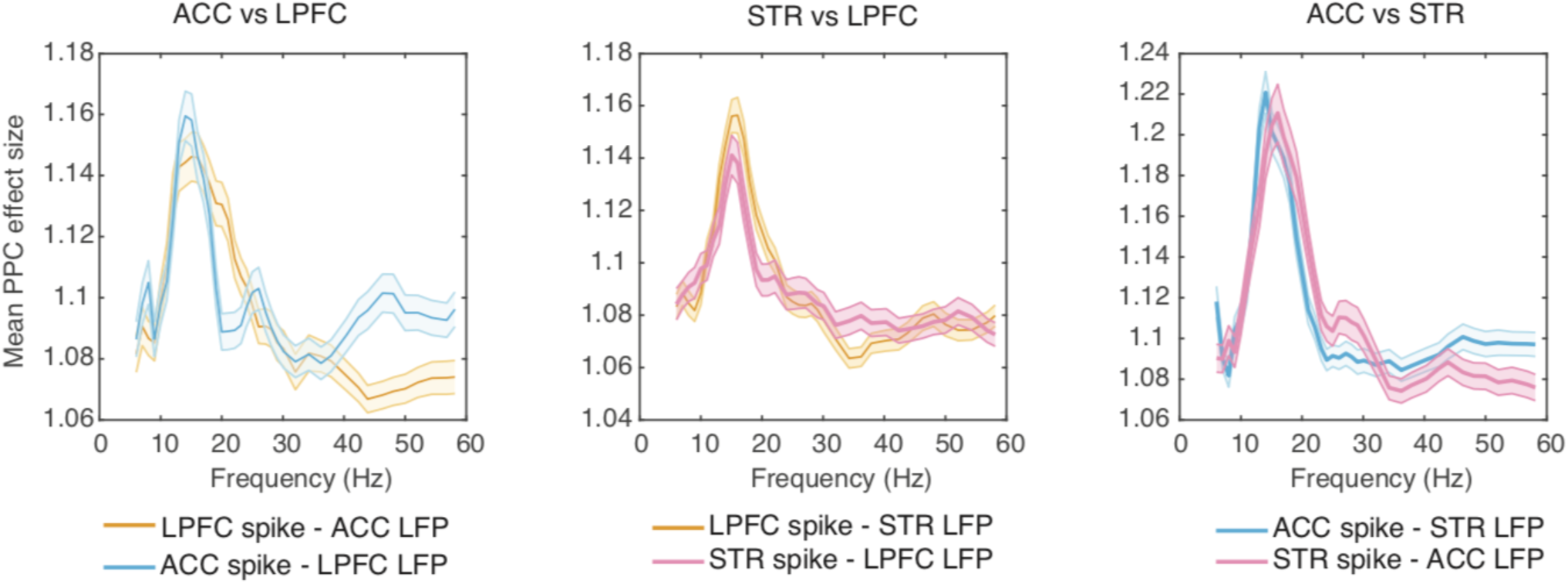
Inter-areal synchronization. Mean and standard deviation of spike-LFP phase synchronization (PPC) between ACC and LPFC (left), STR and LPFC (middle) and between ACC and STR (right).

**Supplemental Figure S3.**
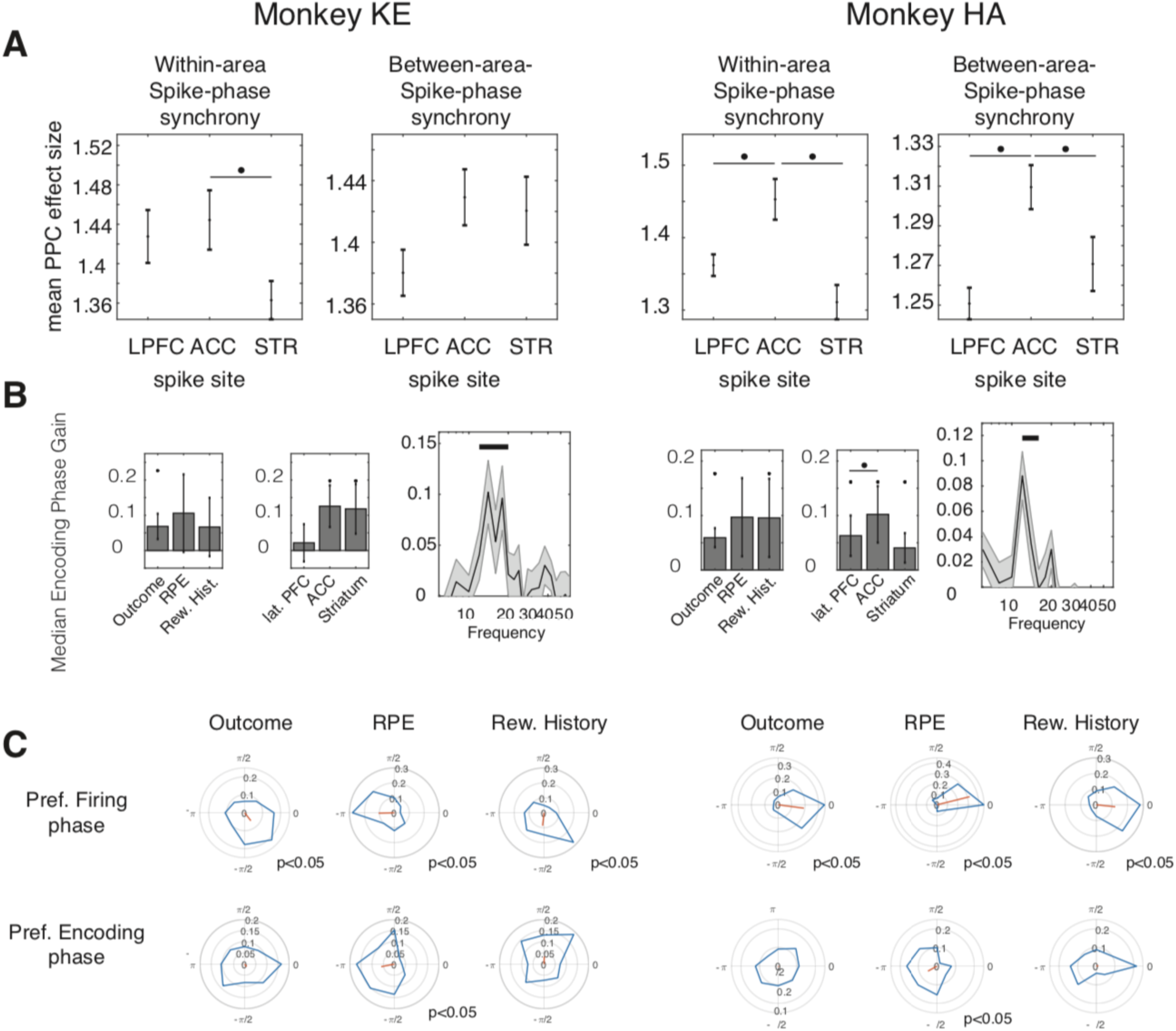
Summary of main results for individual monkeys. Individual results for monkey KE (left) and HA (right). **(A)** Average spike-LFP phase synchronization for spike-LFP pairs within and between areas. (**B**) Average Encoding Phase Gain for each encoding metric (*left*), brain area *(middle*), and across frequencies (*right)* (**C**) Polar histograms of the preferred firing phase (upper panels) and maximal encoding phase (bottom panels) for each encoding metric.

**Supplemental Figure S4.**
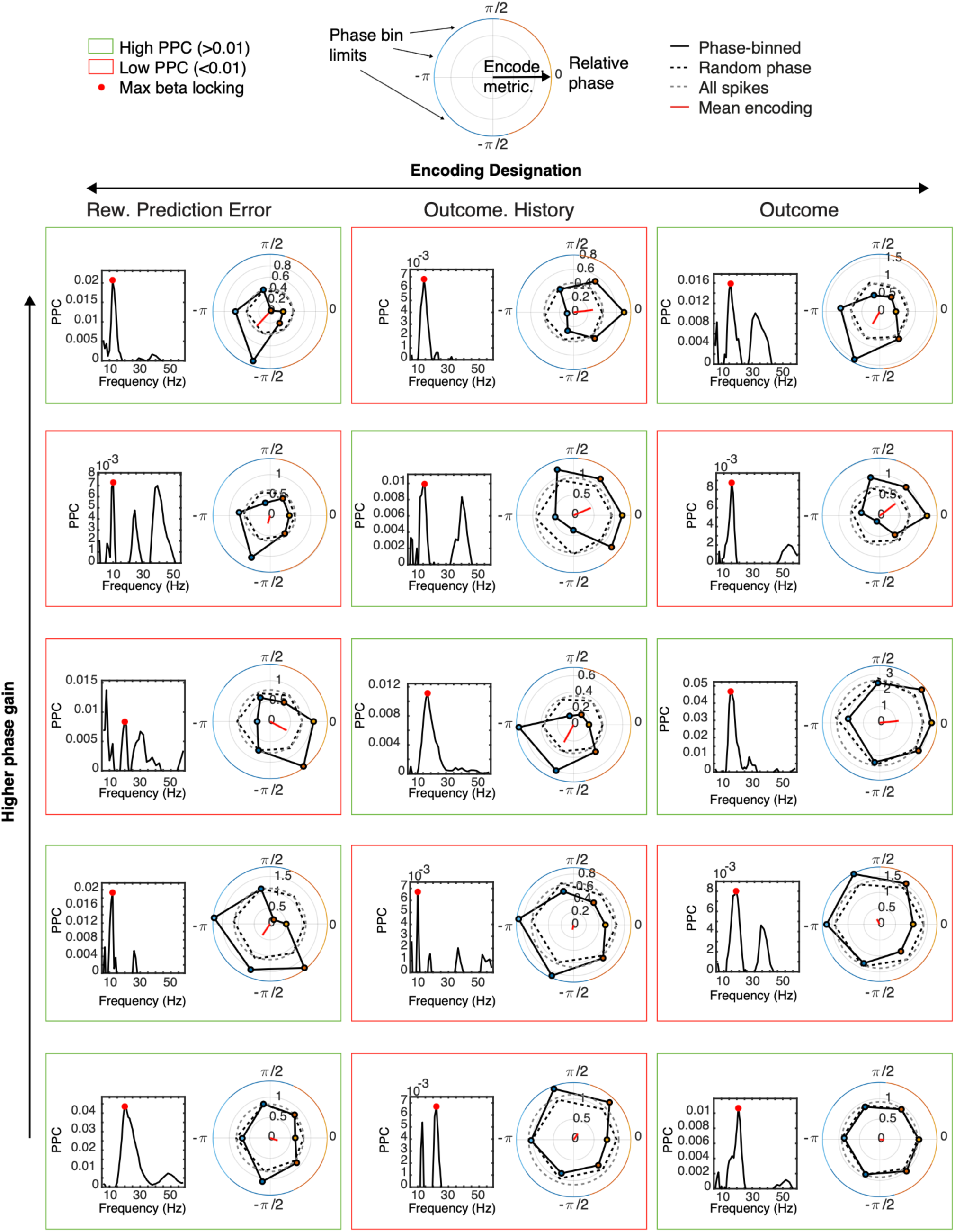
Encoding Phase Gain examples. Columns depict the functional designation. Rows are ordered according to the relative phase gain, with lower phase gain at the bottom, and high phase gain at the top. The spike-phase consistency is depicted on the left, with the maximal significant locking in the [10 25] Hz beta band signified with the red dot. The corresponding phase-dependent encoding is depicted on the right. 0 corresponds to the preferred firing phase. Numbers on concentric circles are the value of the encoding metric. The grey dotted line represents the encoding metric estimated using all spikes, whereas the black dotted line represents the average across many spike-phase randomizations. The red line is the average direction. Colored border lines represent size of the bin. The colored box represents examples with a high (green) or low (red) degree of synchrony.

**Supplemental Figure S5.**
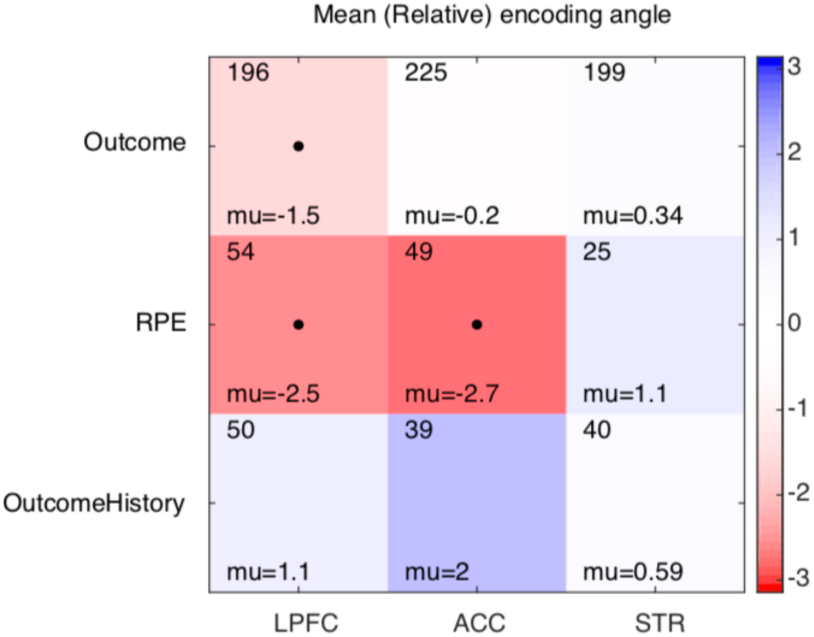
Preferred encoding for each area and function. Mean preferred encoding phase for *Outcome, RPE*, and *Outcome History* (y-axis) cells in the ACC, LPFC, and STR (x-axis). Color represents the (relative) encoding. Black dots represent significant phase concentration (Hodge-Ajne test, p<0.05). The number of spike-LFP pairs that went into each cell is depicted in the top left, and the mean phase is on the bottom left.

